# ASCT: automatic single-cell toolbox in julia

**DOI:** 10.1101/2023.12.27.573479

**Authors:** Ling Yang, Nan Li

## Abstract

ASCT(https://github.com/kaji331/ASCT) is an automatic single-cell toolbox for analyzing single-cell RNA-Seq data. This toolbox can analyze the output data of 10X Cellranger for quality checking, preprocessing, dimensional reduction, clustering, marker genes identification and samples integration. It completely runs all functions by automatic methods without artificial intervention and can tune the parameters for advanced user. It is implemented by pure Julia language, and the overall runtime of basic steps is less than Seurat V4.

## Background

There are many frameworks for single-cell RNA-Seq analysis, such as Seurat[1], Scanpy[2] and etc, were built by R/Python languages. Some of these frameworks need C/C++ or other static language background for acceleration, and some of these frameworks need user manually select parameters using diagnostic plotting or external tools, like PCs, cluster numbers etc. Here, we present a framework that only use Julia language for high performance and simple development. Moreover, in contrast to the existing frameworks, ASCT minimizes the manual tuning steps as much as possible, and basically provides a foolproof operation for beginner-level users while adapting to different data scenarios in a targeted manner to achieve better analysis results.

## Results

### ASCT workflow

ASCT implements the classical workflow of single-cell RNA-Seq analysis established by previous frameworks with extra optimizations, including data fetching of 10X Cellranger output, quality control by clustering and linear regression methods, data preprocessing, dimensional reductions by PCA/UMAP[3]/tSNE[4,5], clustering using the Louvain algorithm[6], marker genes identification by hypothesis testing and COSG algorithm[7] (Fig.1), samples integration using the Harmony algorithm[8] etc.

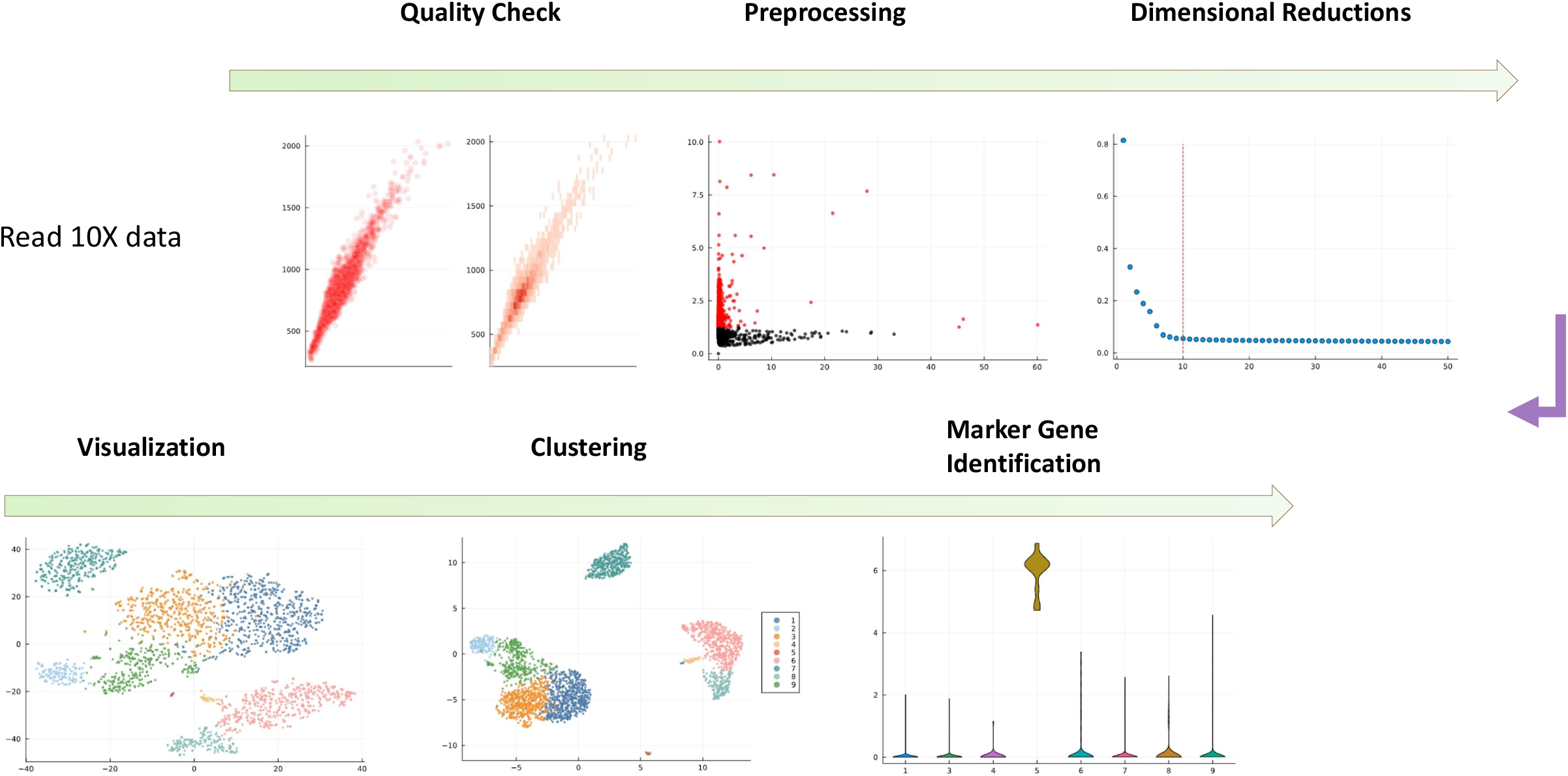

### Compare ASCT with other packages

We used the pbmc3k data set for benchmarks to Seurat V4 (https://github.com/satijalab/seurat/blob/4d185f8d1a2d3f187904b89ed7c0bd47e2bc765f/vignettes/pbmc3k_tutorial.Rmd). All steps did not need diagnostic plotting and parameter tuning, offering very similar results and shorter processing time for the entire workflow (Fig. 2A) even including extra computational processes (https://github.com/kaji331/ASCT/blob/main/doc/pbmc3k.ipynb). Moreover, we used the neuron9k data set for more run-time benchmarks (https://github.com/kaji331/ASCT/blob/main/doc/pbmc3k.ipynb) showing better results by ASCT’s automatic parameters (Fig. 2B). Thus, ASCT provides a fast pipeline for interactive or automatic analysis of single-cell RNA-Seq data sets.

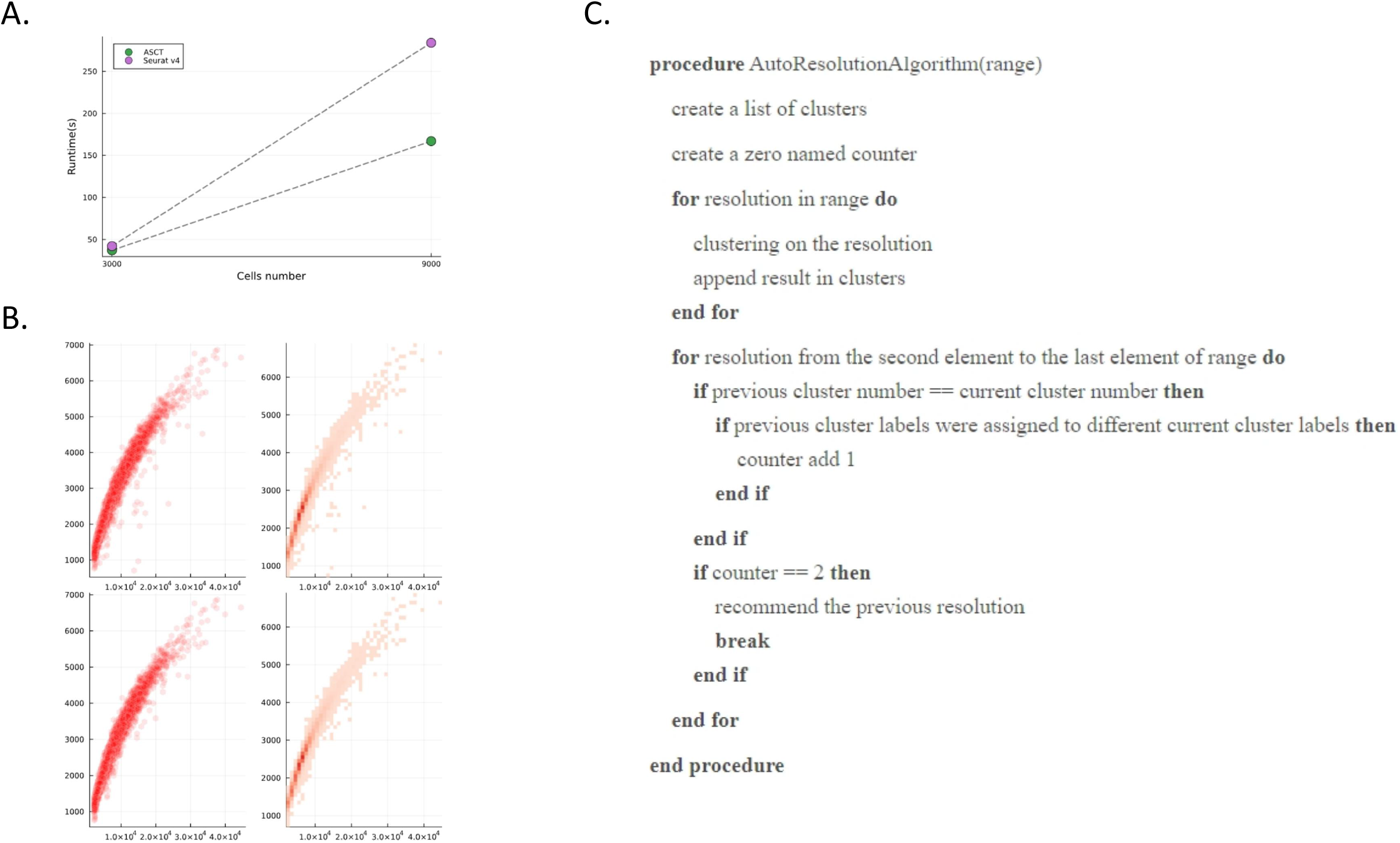

For Louvain algorithm, referring to the thought of clustree[9] package, we implemented a method to select a common resolution value (Fig. 2C). In addition to the community-detection algorithm for clustering, we also provide automatic k-medoids[10] algorithm for the exploration of different approaches by the silhouettes coefficients and the elbow point of total costs. We also implemented the Harmony algorithm for data integration with satisfactory effect and faster speed than the CCA algorithm (https://github.com/kaji331/ASCT/blob/main/doc/Harmony.ipynb).

### Conclusions

ASCT’s performance meets the requirements for most common data analysis of single-cell RNA-Seq, while its fully automatic process greatly reduces the difficulty for novice users and probably helps minimize the generation of unreasonable results. Furthermore, ASCT could be conveniently developed solely using the Julia programming language. With the continuous evolution of Julia language and the expansion of its community, the functionality of ASCT will continue to increase, and the performance will be further enhanced. The data exchange to other packages like Seurat or Scanpy would be implemented easily by HDF5 file format, which is language and platform independent. The construction of ASCT relied on the abundant resources in the existing Julia computing community, while also expanding the ecosystem of bioinformatics in Julia.

## Methods

### ASCT’s fundamental dependencies

ASCT currently relies primarily on HDF5[11], SparseArrays[12], DataFrames[13] etc. The dimensional reductions need MultivariateStats[14], UMAP and TSne. The clustering relies on Distances[15], LinearAlgebra[16], Graphs[17], NearestNeighbors[18]. The marker genes identification relies on HypothesisTests[19] and MultipleTesting[20]. Some implementations of parameters selection need GLM[21], Loess[22], Clustering and KernelDensity[23].

Julia’s machine-learning packages, like Flux, could enrich the application of ASCT. However, compared to PyTorch/TensorFlow in Python, these packages need broader support from developers and further optimization. For classical statistical algorithms, Julia still requires more effort in terms of advanced methods and expanding implementations.

### Comparison with existing frameworks

Now, Seurat in R and Scanpy in Python are dominating the field of single-cell analysis applications, even have expanded into the realm of multimodal analysis. In contrast to CellScopes[24] in Julia, The ASCT is just a beginning for single-cell RNA-Seq analysis, no modules for ATAC and Spatial data, but focusing primarily on foolproof adaptive analysis and a certain extent performance optimization, rather than just being a repetitive substitute for Seurat/Scanpy. The excellent BPCells[25] in R announced a disk-backed method for atlas-level analysis of single-cell RNA-Seq/ATAC-Seq data sets with impressive high speed and low memory requirement. Optimizing the ASCT in a manner like BPCells while maintaining the capability for automatic analysis would be a milestone for future development.

## Availability and requirements

ASCT’s open-source code are maintained on Github (https://github.com/kaji331/ASCT) and published under the BSD-3 license. All demonstrations and benchmarks in the main text are stored at https://github.com/kaji331/ASCT.

The two used data sets were downloaded from the website of 10X Genomics.

The developing programming language: Julia v1.6.7

## Acknowledgments

We thank the authors of Seurat, Scanpy, BPCells, CellScopes and Clustree for sharing their great tutorials. We are also grateful to the community of Julia language. The High-Performance Computing Center of Westlake University supported the whole development and testing of ASCT.

## Ethics declarations

Ethics approval and consent to participate

Ethics approval was not applicable for this study.

